# Dendro-somatic synaptic inputs to ganglion cells violate receptive field and connectivity rules in the mammalian retina

**DOI:** 10.1101/2021.07.01.450751

**Authors:** Miloslav Sedlacek, William N. Grimes, Morgan Musgrove, Amurta Nath, Hua Tian, Mrinalini Hoon, Fred Rieke, Joshua H. Singer, Jeffrey S. Diamond

**Author notes:** These authors contributed equally (Miloslav Sedlacek, William N. Grimes).

## Abstract

In retinal neurons, morphology strongly influences visual response features. Ganglion cell (GC) dendrites ramify in distinct strata of the inner plexiform layer (IPL) so that GCs responding to light increments (ON) or decrements (OFF) receive appropriate excitatory inputs. This vertical stratification prescribes response polarity and ensures consistent connectivity between cell types, whereas the lateral extent of GC dendritic arbors typically dictates receptive field (RF) size. Here, we identify circuitry in mouse retina that contradicts these conventions. A2 amacrine cells are interneurons understood to mediate “cross-over” inhibition by relaying excitatory input from the ON layer to inhibitory outputs in the OFF layer. Ultrastructural and physiological analyses show, however, that some A2s deliver powerful inhibition to OFF GC somas and proximal dendrites in the ON layer, rendering their inhibitory RFs smaller than their dendritic arbors. This OFF pathway, avoiding entirely the OFF region of the IPL, challenges several tenets of retinal circuitry.

## Introduction

AII (A2) amacrine cells (ACs) play diverse signaling roles in the mammalian retina (Demb and Singer, 2012; Marc et al., 2014). During night (scotopic) vision, A2s relay input from rod bipolar (RB) cells to ON and OFF cone bipolar (CB) cell axon terminals that contact GCs (Figure 1A). ON signals from RBs are preserved via electrical synapses to ON CBs and inverted via inhibitory, glycinergic synapses to OFF CBs (Famiglietti and Kolb, 1975; Pourcho and Goebel, 1985; Strettoi et al., 1992; Tsukamoto et al., 2001; Figure 1A). This circuitry reliably transmits scotopic signals to ON GCs (Smeds et al., 2019), but A2 inhibition of OFF CBs exerts little influence on OFF GC signaling near visual threshold (Arman and Sampath, 2012). A2s also contact OFF GCs directly (Manookin et al., 2008; Murphy and Rieke, 2008; Münch et al., 2009; van Wyk et al., 2009; Beaudoin et al., 2019), but synaptic connections between individual A2-GC pairs in the OFF layer of the IPL are weak (Münch et al., 2009; Graydon et al., 2018). It therefore remains unclear how visual signals are transmitted most reliably to OFF GCs at night.

**Figure 1.**
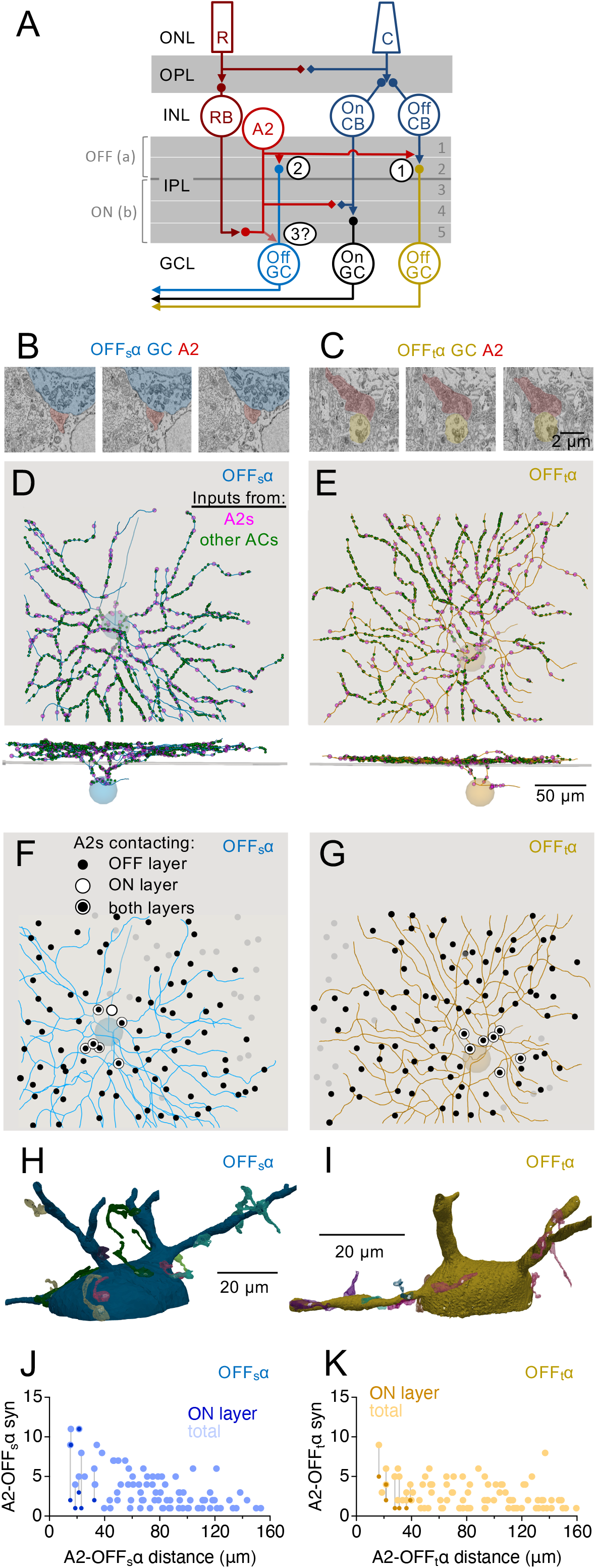
A2 ACs make chemical synapses onto OFFα GC somas. **(A)** Retina schematic (modified from Vaney et al., 1991). ◆◆, gap junctions; ▶●, chemical synapses. *1*, A2 synapses onto the axon terminals of OFF CBs; *2*, A2 synapses onto OFF GC dendrites; *3*, proposed output synapses onto the somatic region of OFF GCs. R, rod photoreceptor; C, cone photoreceptor; RB, rod bipolar cell, A2, A2 amacrine cell; On CB, ON cone bipolar cell; Off CB, OFF cone bipolar cell; Off G, OFF ganglion cell; On G, ON ganglion cell; ONL, outer nuclear layer; OPL, outer plexiform layer; INL, inner nuclear layer; IPL, inner plexiform layer; GCL, ganglion cell layer. **(B)** Serial SBFSEM micrographs showing a synapse from an A2 AC onto the soma of an OFF_s_α GC. Series shows every other acquired section, therefore depicting 52-nm intervals. **(C)** As in **(B)**, but showing an A2 synapse onto a proximal dendrite of an OFF_t_α GC. Scale bar applies to **(B,C)**. **(D-E)** Skeletonized OFF_s_α **(D)** and OFF_t_α **(E)** GCs with all inhibitory inputs annotated. Lower panels show side (transverse) views. Gray plane in transverse views indicates the raminfication of OFF starburst ACs (SACs). Scale bar in **(E)** applies to all panels in **(D-G)**. See Figure S1 for annotated excitatory inputs. **(F-G)** Soma locations of A2s providing inputs to OFF_s_α **(F)** and OFF_t_α **(G)** GCs in the OFF (black circles) and/or ON layers (white circles) of the IPL. Gray circles indicate unconnected A2s. See Figure S2 for inputs from other ACs. **(H-I)** 3D images of OFF_s_α **(H)** and OFF_t_α **(I)** GC somatic region. Presynaptic A2s are rendered in different colors; chemical synapses are white. **(J-K)** Summary plots showing the total number of chemical synaptic contacts from A2s to the OFF_s_α **(J)** and OFF_t_α **(K)**, and those in the ON layer of the IPL, as a function of retinotopic (*x-y*) distance between the A2 and GC somas.

Here, we present ultrastructural analysis of mouse retina indicating that that A2s make chemical synapses deep in the IPL and ganglion cell layer (GCL) onto the proximal dendrites, soma, and axon initial segments of sustained and transient OFFα (OFF_s_α and OFF_t_α) GCs. These synapses are made by a small subset of A2s whose distal, arboreal dendrites encounter OFFα GC somas. The synaptic outputs arise from the distal tips of the A2 arboreal dendrites, close to where A2s receive excitatory inputs from RBs, ensuring efficient signal transfer from input to output. Fluorescence imaging and electrophysiology experiments indicate that depolarizing stimuli elevate [Ca^2+^] in A2 arboreal dendritic compartments and elicit glycinergic inhibitory postsynaptic currents (IPSCs) in OFFα GCs. Scotopic light-evoked IPSCs in OFF_s_α GCs exhibit narrow RFs that do not correlate with dendritic arbor dimensions. Together, these results show that OFFα GCs receive powerful somatic inhibition that influences RF properties and that A2 synaptic organization is heterogeneous and depends on proximity to OFFα GC somas.

## Results

In the canonical view of mammalian retinal circuitry (Demb and Singer, 2012), A2s make all their glycinergic outputs onto CBC axon terminals and GC dendrites from their lobular appendages in sublamina a (the “OFF layer”) of the IPL (Famiglietti and Kolb, 1975; Manookin et al., 2008; Murphy and Rieke, 2008; Münch et al., 2009; van Wyk et al., 2009; Beaudoin et al., 2019; Figure 1A). A2s also extend arboreal dendrites deep into sublamina b (the “ON layer”) of the IPL, where they receive excitatory inputs from RBs and make electrical synapses onto ON CBC axon terminals (Famiglietti and Kolb, 1975; Pourcho and Goebel, 1985; Dacheux and Raviola, 1986; Strettoi et al., 1992; Mills and Massey, 1995) but make chemical synaptic outputs only rarely (Tsukamoto and Omi, 2013). A2 arboreal dendrites occasionally extend into the ganglion cell layer (GCL; Zandt et al., 2017), but their functional roles have not been examined. Below, we describe anatomical and physiological analyses that identify and investigate synaptic connections between A2s and the somatic regions of OFFα GCs.

### EM reconstructions reveal selective connections between A2s and OFFα GCs in the ON layer

While examining A2 ultrastructure and synaptic circuitry within a conventionally stained serial block-face scanning electron microscopy (SBFSEM) volume from mouse retina (Ding et al., 2016; Graydon et al., 2018; Park et al., 2020), we observed that some A2s contained presynaptic active zones within their arboreal (i.e., ON-layer) dendrites (e.g., Figure 1B,C). As these active zones appeared presynaptic to GC dendrites and somas, we systematically annotated “ON/GCL” synaptic connectivity between A2s and GCs. Beginning at presynaptic structures within the arboreal dendrites of previously identified and traced (“skeletonized”) A2s (Graydon et al., 2018; Park et al., 2020), we followed postsynaptic GC dendrites and completely skeletonized two GCs that appeared, based on dendritic stratification and soma size, to be OFF_s_α and OFF_t_α GCs (Figure 1D,E). One of each OFFα GC was found within the data set, consistent with OFF_s_α and OFF_t_α GC density in rat retina (Roy et al., 2021). Notably, no other GCs were found to be postsynaptic to ON/GCL synapses made by these A2s.

We identified and annotated all of the excitatory (Figure S1) and inhibitory synaptic inputs onto these OFF_s_α and OFF_t_α GCs and found that they receive 1230 and 1097 inhibitory inputs, respectively (Figure 1D,E). We then skeletonized the presynaptic process at each inhibitory input sufficiently to determine whether it arose from an A2 (see STAR Methods for A2 identification criteria) or another AC type. The dendritic arbor of each presynaptic A2 was skeletonized suffiently to identify all of its synaptic contacts with each OFFα GC. The OFF_s_α GC received 278 (22.6%) of its inhibitory inputs from 85 different A2s (Figure 1D,F); the OFF_t_α GC received 253 (23.1%) of its inhibitory inputs from 93 different A2s (Figure 1E,G).

A small fraction (5.8% and 2.5%) of the total inhibitory inputs onto the OFF_s_α and OFF_t_α GC, respectively, were made onto the somas and axon initial segments, as well as proximal dendrites passing through the ON layer (Figure 1D,E). A2s contributed disproportionately to this subset, providing 29 of 72 (40.3%, from 7 different A2s) ON/GCL inputs to the OFF_s_α GC and 16 of 28 (57.1%, from 7 different A2s) to the OFF_t_α GC (Figure 1D-I). Other ON/GCL inputs were provided by wide-field ACs (Figure S2). Notably, A2s that did not provide ON/GCL input to either OFFα GC did not contain presynaptic active zones in their arboreal dendrites, even in dendrites closely apposed to somas and proximal dendrites of other GCs. A2 chemical synaptic ON/GCL outputs appear, therefore, to target OFFα GCs exclusively.

Only 11% of A2 OFF-layer outputs were made onto GC dendrites (Graydon et al., 2018), but this minority also targeted OFFα GCs preferentially: of the 7.5 ± 2.7 OFF-layer outputs to GCs from each of 31 completely skeletonized A2s in this data set (including 12 from Graydon et al., 2018), 3.1 ± 2.1 (42 ± 24%) and 2.0 ± 1.5 (27 ± 23%) were made onto the reconstructed OFF_s_α and OFF_t_α, respectively. As each point in retinal space is covered by roughly 2 OFF_s_α and 2 OFF_t_α GCs (Bae et al., 2018), many of the remaining outputs likely targeted dendrites of OFFα GCs whose somas were not contained in the data set. These results corroborate recent physiological observations in guinea pig retina that A2s contact primarily GC types corresponding to the OFF_s_α and OFF_t_α GCs described here (Beaudoin et al., 2019).

ON/GCL inputs to OFFα GCs came only from those A2s with somas located almost directly above the GC somas (Figure 1F,G,J,K). Consequently, although many A2s made OFF-layer synapses onto both OFFα GCs, none made ON/GCL inputs to both (Figure 1F,G). The number of ON/GCL inputs from individual A2s varied, ranging from 1 to 11 (4.1 ± 4.1, n = 7) for the OFF_s_α GC and 1 to 5 (2.3 ± 1.6, n = 7) for the OFF_t_α GC; Figure 1H-K). Although we did not reconstruct completely every A2 in the data set, every ON/GCL A2 input to the OFFα GCs could be traced back to a fully reconstructed A2; A2s making ON/GCL synapses also provided OFF-layer input to the same OFFα GC in all but one case (3.6 ± 2.6 synapses onto the OFF_s_α GC and 2.9 ± 1.1 onto the OFF_t_α GC). A2s that failed to make ON/GCL synapses exhibited no apparent differences compared to those that did, making similar numbers of OFF-layer synapses to the OFFα GCs (2.9 ± 2.0 synapses onto the OFF_s_α, 2.5 ± 1.6 onto the OFF_t_α; Figure 1J,K). These similarities suggest that A2s providing ON-layer inputs to OFFα GCs do not represent a distinct A2 subtype (see Discussion); rather, these ON/GCL synapses may arise based on proximity of an A2’s arboreal dendrites to the somatic region of the OFFα GCs. Although ON/GCL A2 inputs constituted only 2.4% and 1.5% of the total number of inhibitory inputs to the OFF_s_α and OFF_t_α GCs, respectively, experiments described below indicated that they contribute substantially to OFFα visual signaling.

### OFFα GCs express glycine receptors in the ON IPL and GCL

We next examined glycine receptor (GlyR) expression in OFFα GCs using fluorescence immunohistochemistry (Figure 2). GlyRs require α subunits to function (Betz and Laube, 2006), and OFFα GCs express α_1_ exclusively (Majumdar et al., 2007). Here, we labeled all GCs with an antibody to RBPMS (Figure 2A; Rodriguez et al., 2014) and observed somatic GlyRα_1_ immunoreactivity (Figure 2B) particularly in a subset of GCs that also expressed SMI-32 (Figure 2C), a marker for α GCs (Lin et al., 2004; Figure 2D).

**Figure 2.**
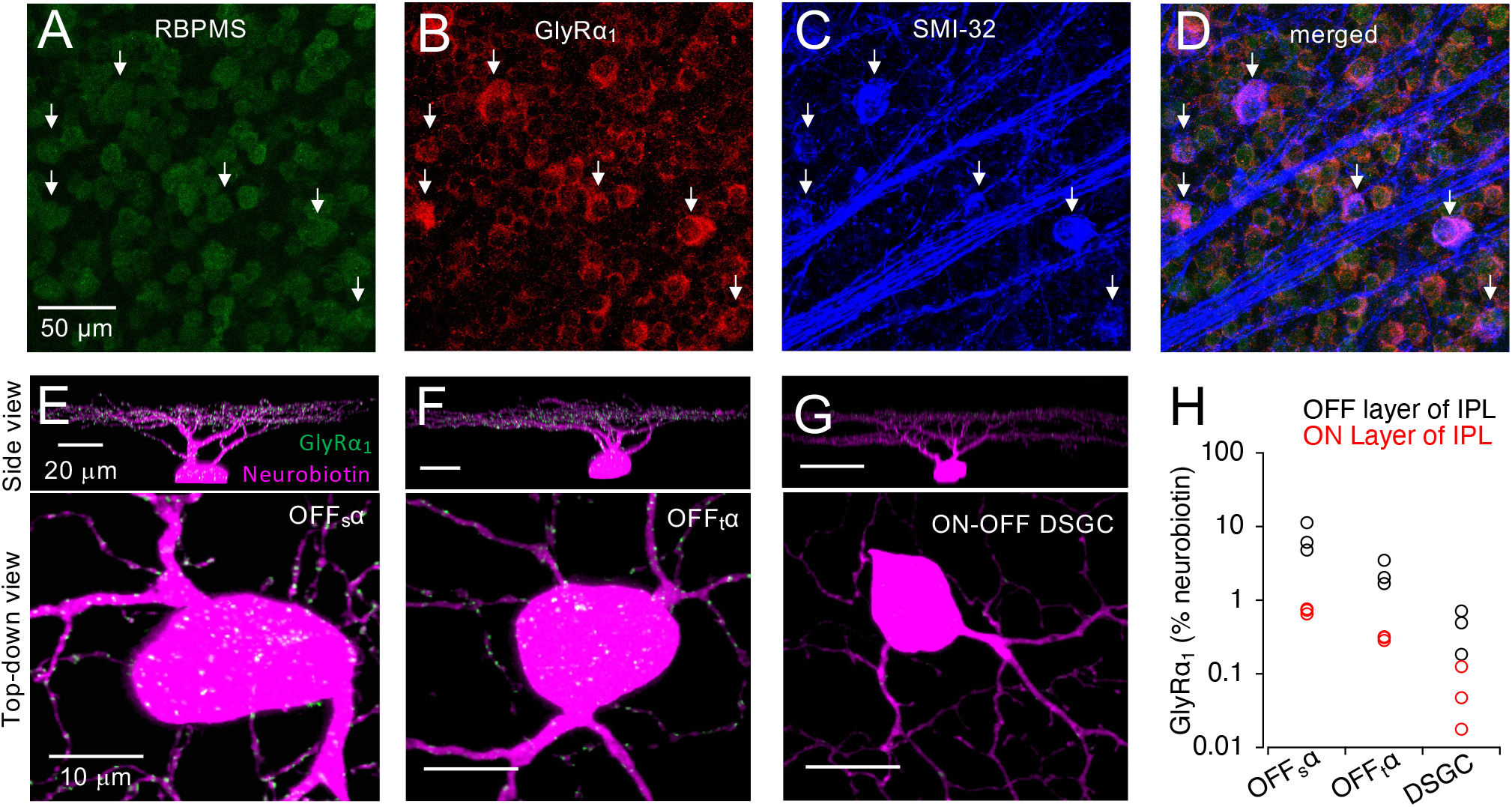
OFFα GCs express GlyRs densely in their somatic region. **(A-D)** Fluorescence micrographs of the GCL of a WT mouse retina incubated in antibodies to the GC marker RBPMS **(A)**, GlyRα_1_ **(B)** the αGC marker SMI-32 **(C)** and a merged image **(D)**. White arrows point to the same αGC somas in each panel. Scale bar in **(A)** applies to **(A-D)**. **(E-G)** Side (*top*) and top-down (*bottom*) views of three neurobiotin (magenta) filled GCs (OFF_s_α **(E)**, OFF_t_α **(F)** and ON-OFF DS **(G))** showing the GlyRα_1_ signal (green) contained within the GC soma and proximal dendrites. **(H)** Quantification of the % occupancy of GlyRα_1_ within the ON and OFF arbors of the three GC types. The soma and the proximal dendritic arbor were included in the ‘ON’ arbor quantification. (n = 3 GCs of each type; n = 3 animals).

To examine GlyR expression specifically in OFFα GCs, individual cells were filled with neurobiotin, then fixed and labeled with antibodies to neurobiotin and GlyRα_1_ (Figure 2E-H). As expected, OFFα GCs robustly expressed GlyRα_1_ in the OFF layer of the IPL (Figure 2E,F): OFF_s_α and OFF_t_α GCs exhibited greater GlyRα_1_ expression in the OFF layer compared to, for example, ON-OFF direction-selective GCs (DSGCs; n = 3 cells of each type, *p* = .02, Mann-Whitney test; Figure 2G), which express primarily GlyRα_2_ and GlyRα_4_ (Pyle, 2019). OFF_s_α and OFF_t_α GCs also exhibited stronger GlyRα_1_ expression than DSGCs in the ON layer (*p* = .02, Mann-Whitney test; Figure 2H), even though DSGC dendrites ramified extensively in this layer (Figure 2G). GlyRα_1_ expression was higher in OFF_s_α GCs than in OFF_t_α GCs in both layers (*p* = .02, Mann-Whitney test; Figure 2H).

### Close apposition of RB inputs and arboreal outputs suggests local input-output relationship

Individual AC types have evolved different strategies to couple synaptic input to output (Diamond, 2017). Some wide-field amacrine cells use active conductances to relay synaptic input signals to outputs located hundreds of microns away (Dacey, 1989; Bloomfield and Völgyi, 2007; Greschner et al., 2014; Manookin et al., 2015). Conversely, A17 ACs couple input to outputs that are 1000 times closer, using Ca^2+^ influx through postsynaptic glutamate receptors to elicit transmitter release ≤100 nm away (Chávez et al., 2006; Grimes et al., 2015). With chemical outputs on both arboreal and lobular dendrites, A2s might integrate inputs on multiple spatial scales to provide output to the same OFFα GCs.

To examine the spatial relationship between inputs and output on A2 arboreal dendrites, we annotated all RB inputs (n = 139) onto A2 terminal arboreal dendrites that contacted the OFF_t_α and OFF_s_α GCs in the EM dataset (Figure 3). All chemical synaptic outputs from A2 arboreal dendrites occurred in the ON layer, within the innermost 42% of the IPL, or in the GCL (Figure 3A). Most RB inputs were made onto the last 20 μm of the A2 terminal dendrite, whereas outputs to OFFα GCs arose primarily within the last 10 μm (Figure 3B). Only one output synapse was located within 300 nm of an RB input, however, arguing against hyperlocal input-output coupling like that observed in A17s.

**Figure 3.**
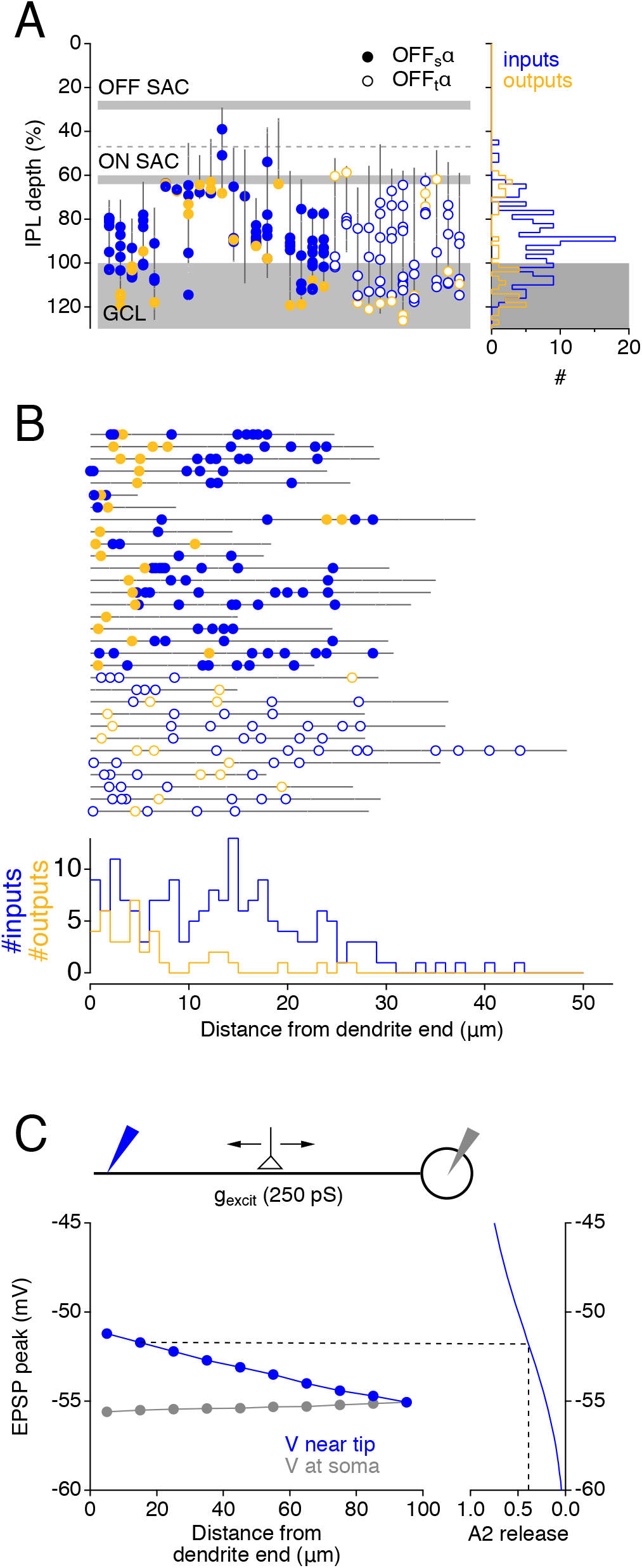
Localization of RB inputs and chemical synaptic outputs in A2 arboreal dendrites. **(A)** Plot showing the IPL depth of 32 A2 terminal dendrites making ON/GCL synapses onto the OFF_s_α (closed gold circles) and OFF_t_α (open gold circles). For clarity, dendrites are rendered as perfectly vertical lines. Inputs from RBs indicated in blue. *Right*, histogram showing IPL depth of RB inputs and outputs to OFFα GCs. **(B)** RB inputs and A2 outputs shown as in (A), except that A2 terminal dendrites have been straightened and aligned by their ends to compare linear distance along each dendrite between RB inputs (blue) and A2 outputs to OFFα GCs (gold). *Bottom*, histogram showing locations of inputs and outputs relative to the dendrite terminus. **(C)** Electronic model comparing simulated EPSPs, recorded near the end of the dendrite (blue) or at the soma (gray), evoked by a 250-pS excitatory synaptic conductance at varying locations along the dendrite. *Right*, Simulated EPSPs projected onto an A2 release function derived from paired recordings between A2s and type 2 CBs (Graydon et al., 2018).

While >300 nm may be too far to permit direct (Ca^2+^-mediated) input-output coupling, their relatively close apposition suggests that excitatory inputs from RBs may depolarize the dendritic membrane sufficiently to activate Ca_v_ channels and elicit release from nearby presynaptic active zones. In particular, dendritic tips are effectively depolarized by even small excitatory synaptic inputs due to the reduced intracellular axial pathways for charge flow (Rall, 1969). [Gap junctions in arboreal dendrites may offer alternative conduction pathways but most are located closer to the soma (Strettoi et al., 1992; Marc et al., 2014)]. A simple electrotonic model, together with the experimentally measured relationship between A2 membrane potential and glycine release (Graydon et al., 2018), illustrates this idea (Figure 3C): An excitatory conductance in the A2 approximating that of a miniature EPSC (Singer et al., 2004; Jarsky et al., 2010) is predicted to depolarize the dendrite sufficiently to evoke release at at nearby output synapses, particularly when the input arises within 20 μm of the sealed end (Figure 3C).

### L-type Ca_v_ channels mediate Ca^2+^ influx into A2 arboreal dendrites

Glycine release from A2 arboreal dendrites likely requires local expression of functional Ca_v_ channels. Mouse A2s express Ca_v_1.3 (L-type) channels, but widefield fluorescence Ca^2+^ imaging methods detected signals primarily in proximal, lobular dendrites (Habermann et al., 2003; Balakrishnan et al., 2015). Arboreal dendrites are thinner and more diffuse, however, possibly making Ca^2+^ signals more difficult to detect. To re-visit this issue, we recorded from A2s in whole-mount mouse retina, filled them with Alexa 594 (50 μM) and the Ca^2+^ indicator dye Fluo-5F (150 μM), and imaged dendrites with confocal microscopy (Figure 4). Visual responses were blocked pharmacologically (see STAR Methods). Depolarizations applied under voltage clamp elicited Ca^2+^ indicator signals in arboreal and lobular dendrites (Figure 4A,B). Indicator ΔF/F_o_ signals increased with the amplitude of the step depolarization, reflecting the voltage dependence of A2 L-type currents (Habermann et al., 2003; Figure 4A,B). Dendritic ΔF/F_o_ signals of varying size exhibited similar voltage-dependence (Figure 4A, *right*), suggesting that responses were not distorted by indicator saturation. Step-evoked indicator signals in all dendrites were blocked by the Ca_v_1 channel antagonist isradipine (10 μM; Figure 4C,D).

**Figure 4.**
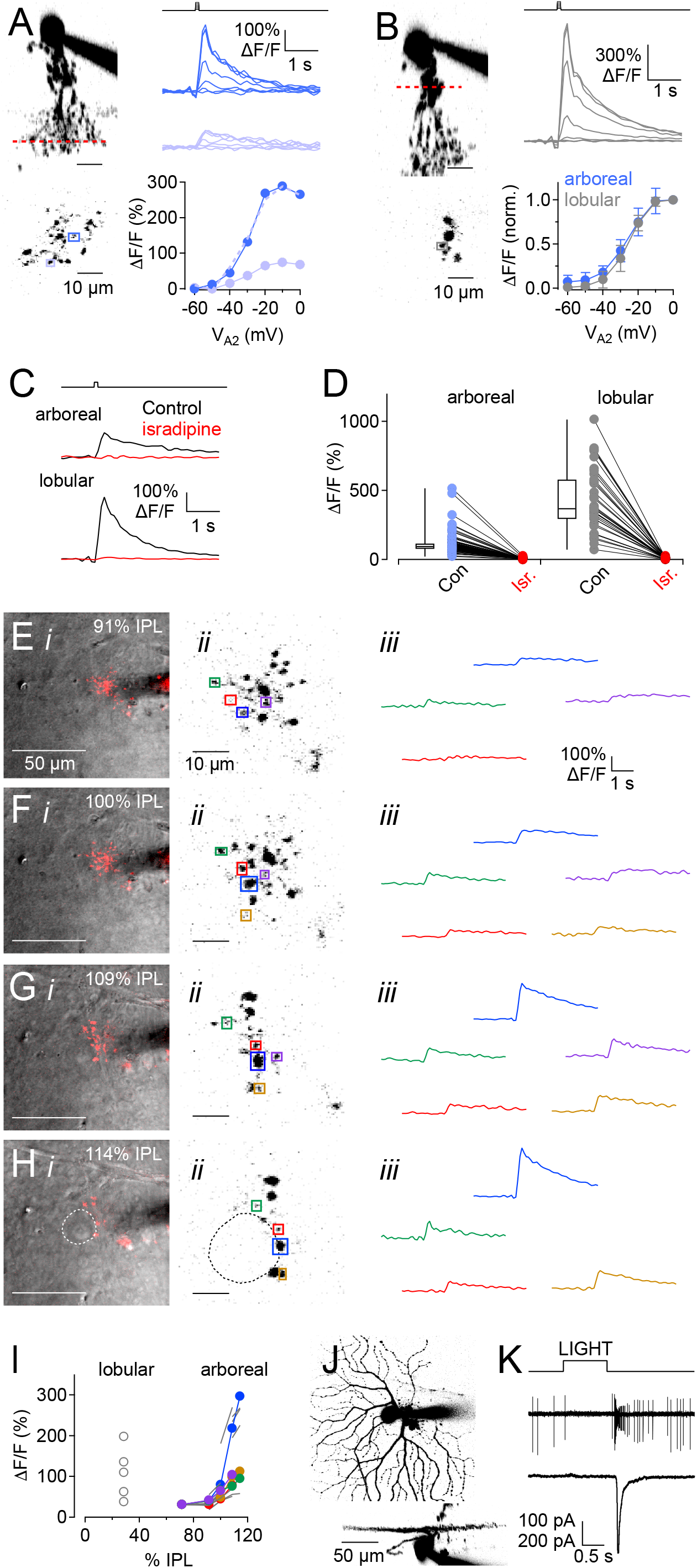
Ca_v_-mediated Ca^2+^ signaling in A2 arboreal dendrites. **(A)** *Top left*, Fluorescence micrograph (*x-z* projection) of a recorded A2 AC filled with Alexa 594 and Fluo 5F. Red line represents focal plane of imaged arboreal varicosities. *Bottom left*, Squares indicate two typical varicosities imaged in a single X-Y optical section. *Top right*, Average indicator ΔF/F signals in varicosities (colors correspond to squares, lower left) evoked by 100-ms voltage steps from −60 to 0 mV (10 mV increments, above). *Bottom right*, Indicator peak ΔF/F as a function of voltage step in the two varicosities. Dashed line show smaller responses scaled to facilitate comparison with the larger responses. **(B)** *Top left*, Fluorescence micrograph of a recorded A2, similar to **(A)** except that the indicator imaging focal plane (red line) was in the lobular region. *Bottom left*, A square highlights a representative example of one lobular appendage in a single X-Y optical section. *Top right*, Average indicator ΔF/F signals in a varicosity (indicated by square in lower left) evoked as in **(A)**. Lower right, voltage dependence of peak indicator ΔF/F signals in arboreal and lobular dendrites. Data in each varicosity were normalized to the ΔF/F signal evoked by a voltage step to 0 mV. **(C)** Average indicator signals in arboreal and lobular dendrites evoked by a step to 0 mV in control conditions (black traces) and in the presence of Ca_v_1 channel antagonist isradipine (10 mM, red traces). Arboreal and lobular regions imaged in two different cells; control vs. isradipine comparisons were made in the same varicosities within each cell. **(D)** Summary showing effects of isradipine on peak indicator signals in arboreal (n = 97) and lobular (n = 33) compartments of A2s. **(E-H)** Ca^2+^ indicator responses imaged in A2 dendrites at four different focal planes in the IPL and GCL. *i*, fluorescence images of the Alexa 594-filled A2 superimposed on a scanning DIC images of the tissue at the indicated depths relative to the top (0%) and bottom (100%) of the IPL, i.e., **(G-H)** were imaged in the GCL. Dashed circle in **(H)** indicates location of OFF_t_α GC soma recorded from in **(J-K)**. *ii*, fluorescence images of the A2 dendrites at each indicated focal plane, with multiple ROIs indicated by squares. Squares of the same color correspond to the same dendrite at different depths. Not all dendrites were recorded at all depths. **(I)** ΔF/F amplitudes as a function of IPL depth recorded from cell shown in **(E-H)**. Each data point represents a single ROI imaged at a particular IPL depth. Colors correspond to examples shown in **(E-H)**. Gray lines indicate ROIs not shown in **(E-H)**. ΔF/F amplitudes from 5 lobular appendages, imaged in the same cell, are shown for comparison (open circles). **(J)** Micrographs (*x-y, top; x-z, bottom*) of an OFF_t_α GC (dashed circle in **(H)**) that closely apposed arboreal dendrites of the recorded A2. After imaging calcium in the A2, the GC was filled with Alexa 594 through a patch electrode and light responses were recorded to identify its morphological and physiological characteristics. Scale bar applies to both views. **(K)** Light responses recorded from the GC shown in **(J)**. Top trace indicates timing of a spot stimulus (300 μm diameter, 100% contrast). Spike responses (*middle*) and EPSCs (*bottom*) were recorded in the cell-attached and whole-cell voltage-clamp configurations, respectively.

Indicator ΔF/F_o_ amplitudes varied widely across neighboring arboreal varicosities (Figure 4A), suggesting that these signals do not reflect passive Ca^2+^ diffusion from more proximal regions of the cell. Accordingly, indicator signals deep in the IPL/GCL were often larger than in more proximal regions of the same dendrites (Figure 4E-I), suggesting that they reflected local Ca^2+^ influx directly into arboreal dendrites. This relative increase in signal was more prevalent in dendrites that extended into the GCL near large GC somas characteristic of α GCs. Responsive arboreal varicosities were often immediately adjacent to α GC somas (e.g., Figure 4H). In several experiments, GC type was confirmed by filling the identified cell with Alexa 594 and recording light-evoked spikes and EPSCs. For example, recordings from the soma visualized in Figure 4H revealed morphological and physiological characteristics of an OFF_t_α GC (Figure 4J,K).

### Strongest A2 inputs come from cells located directly above OFFα GCs

SBFSEM analysis indicated that A2-OFFα ON/GCL connections occur only when the presynaptic A2 is positioned almost directly above the OFFα GC soma (Figure 1). For example, all of the ON layer inputs to the OFF_s_α came from A2s whose somas were located within 33 μm of OFF_s_α in retinotopic space (Figure 1F). By contrast, A2 inputs in the OFF layer were more evenly distributed: A2s located within 36 μm of the OFF_s_α made a similar number of OFF connections (4.7 ± 2.3 synapses, n = 10) to those A2s located 36-72 μm away (3.9 ± 2.2 synapses, n = 25, *p* = .36, *t* test). A2 OFF connections to OFF_t_αs also were evenly distributed across this spatial scale (≤36 μm: 3.4 ± 1.4 synapses, n = 10; 36-72 μm: 2.9 ± 1.5 synapses, n = 25; *p* = .35, *t* test).

These data suggest that A2s located within 36 μm of an OFFα may provide stronger synaptic input to OFFα GCs, provided that ON layer synapses are functional. To test this, we recorded from OFFα GCs and measured IPSCs elicited by activating single presynaptic A2s (Figure 5). Mice expressing Cre recombinase under control of the *Neurod6* promoter (*Neurod6^Cre^*) were crossed with the Ai27 (Gt(ROSA)26Sor) line that expresses channelrhodopsin (ChR2) and tdTomato in a Cre-dependent fashion (Figure 5A). Although the *Neurod6^Cre^* line expresses Cre in a non-A2 narrow field AC (Kay et al., 2011), crossing it with Ai27 yielded ChR2 and tdTomato expression in A2s as well, possibly reflecting leaky expression driven by the ROSA26 cassette (Prabhakar et al., 2019; Figure 5A). The AC types labeled in this line could be distinguished morphologically in *z* stacks obtained at the end of the experiment (Figure 5B; see STAR Methods); only cells identified as A2s were analyzed here.

**Figure 5.**
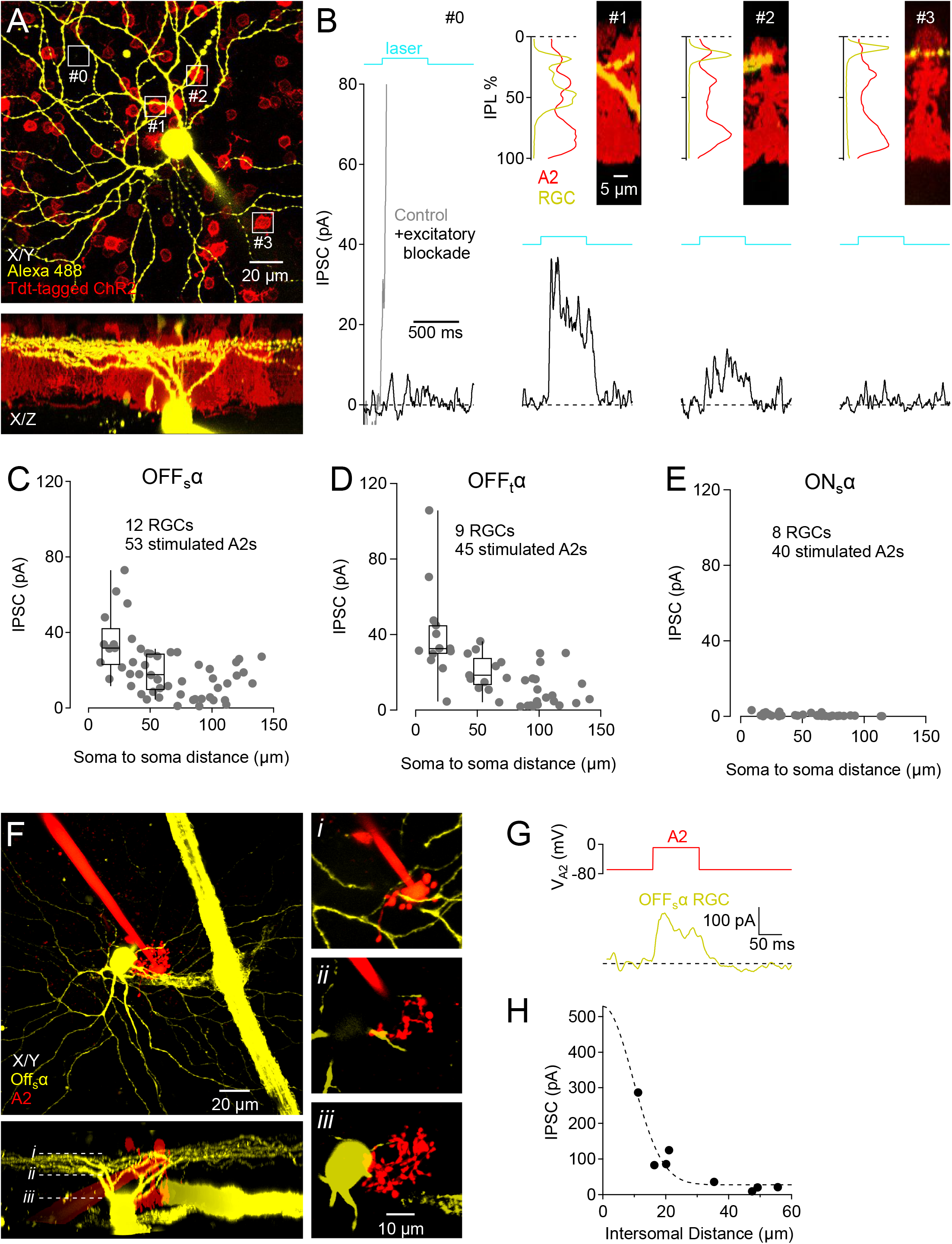
Direct synaptic connections between A2 ACs and OFFα GCs. **(A)** Fluorescence micrographs of an OFF_s_α GC filled with Alexa 488 (yellow) in a retina expressing ChR and tdTomato under control of the NeuroD6 promoter. **(B)** IPSCs (V_hold_ = +10 mV) evoked by laser stimulation directed to the areas indicated by white quares in **(A)**. Respones at location #0 (gray trace, *left*) reflect visual reponses due to photoreceptor stimulation and were eliminated by blockers of BP activation. Additional panels show IPSCs (*bottom*) evoked by stimulating individual A2s, which were identified morphologically by their dendritic ramification pattern in the IPL (*top*). **(C-E)** Summary plots showing IPSC amplitudes recorded in OFF_s_α **(C)**, OFF_t_α **(D)** and ON_s_α **(E)** GCs and evoked by ChR activation of A2s, as a function of distance between the A2 and GC somas. Box plots in **(C)** and **(D)** compare median, range and quartiles of responses to A2s located 0-36 μm and 36-72 μm from the recorded GC. **(F)** Fluorescence micrograph showing a paired recording between a presynaptic A2 (red) and a postsynaptic OFF_s_α GC (yellow). White lines in lower (*x-z*) panel indicate focal planes shown in (*i-iii*). Scale bars apply to left and right panels, respectively. **(G)** Averaged IPSC (V_hold_ = +10 mV) in OFF_s_α GC shown in (F) evoked by step depolarization of the presynaptic A2. **(H)** Summary data from 8 experiments showing the average IPSC amplitude as a function of distance in the *x-y* plane between recorded cells. Dashed line indicates fit by a Gaussian function (λ = 13.25 μm).

ChR2 was activated with 500-ms pulses of a 488-nm laser directed through the microscope to individual A2 somas, and IPSCs (V_hold_ = +10 mV) were recorded from voltage clamped OFFα GCs. Excitatory inputs to ON and OFF bipolar cells were blocked pharmacologically (see STAR Methods), eliminating photoresponses to stimuli delivered in a region lacking tdTomato signal (Figure 5B). ChR-mediated responses were largest when the soma of the stimulated A2 was close (in *x-y* space) to that of the recorded αGC (Figure 5A,B). In OFF_s_α GCs, stimulating A2s located within 36 μm in *x-y* space evoked responses that were almost twice as large (median = 31.8 pA, n = 15) as those evoked by A2s located 36-72 μm away (median = 17.7 pA, n = 16; *p* = .0013, Mann-Whitney test; Figure 5C). Similar results were observed in OFF_t_α GCs (0-36 μm: median = 32.6 pA, n = 13; 36-72 μm: median = 18.5, n = 11; *p* = .0040, Mann-Whitney test; Figure 5D). No significant differences in IPSC amplitudes were observed between OFF_s_αs and OFF_t_αs in either group (0-36 μm: *p* = .65; 36-72 μm: *p* = .72, Mann-Whitney test). A2 stimulation elicited no responses in ON_s_α GCs (Figure 5E), confirming A2 input specificity and successful blockade of bipolar cell activation.

To examine further direct synaptic connections between A2s and OFFα GCs, we obtained whole-cell voltage clamp recordings simultaneously from A2s and OFF_s_α GCs in whole-mount wild-type retina (Figure 5F). Depolarizing voltage steps delivered to the A2 evoked large IPSCs in the OFF_s_α GC when the somas of the two cells were located close to one another in *x-y* space (Figure 5G,H). Together with our anatomical data, these results indicate that A2s positioned above OFFα somas provide stronger inhibitory input, compared with peripherally located A2s, to postsynaptic OFFα GCs.

### ON layer inputs convey rod signals near visual threshold

A2 inputs in the ON layer may render OFF_s_α RGCs particularly sensitive to light stimuli activating just a few A2s. To test this, we made cell-attached recordings from OFF_s_α RGCs and measured spike responses to dim, small (50-μm-diameter spots) light flashes (10 ms; Figure 6A,B). Even stimuli activating just ^~^60 rods (0.071 R*/rod; Jeon et al., 1998), significantly decreased the baseline firing rate (by 38.6 ± 16.6%, n = 7, *p* = .0011, paired *t* test), and a 20× stronger flash eliminated spiking almost completely (95.4 ± 5.3% reduction; Figure 6B).

**Figure 6.**
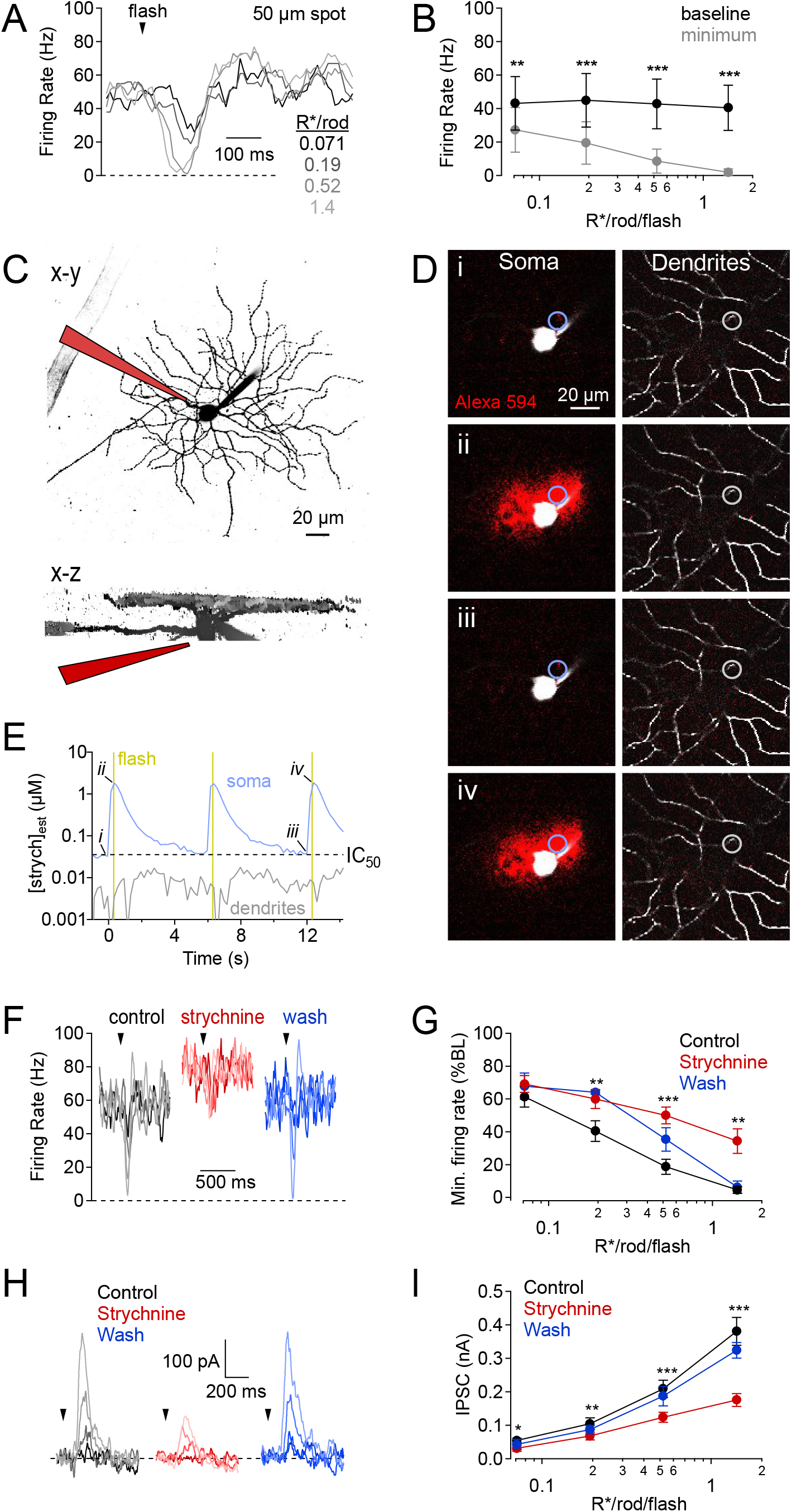
ON/GCL glycinergic inhibition influences OFFα GC light responses. **(A)** Spike frequency plot showing responses in an OFF_s_α GC to brief (10-ms) light flashes of varying intensities. **(B)** Summary data showing baseline and light-evoked minimum firing rates in 7 OFF_s_α GCs. Asterisks indicate paired t-test comparisons between evoked and baseline rates at each intensity. ** = *p* < .01, *** = *p* < .001. **(C)** Fluorescence micrograph of Alexa 488-filed OFF_s_α GC, with approximate location of the strychnine puffer pipette indicated schematically. Scale bar (20 μm) applies to both views. **(D)** Diffusion of Alexa 594 (red) from the puffer pipette, relative to an Alexa 488-filled OFF_s_α (white) at the times indicated in **(E)**. Scale bars in (*i*) apply to lower panels. Circles indicate ROIs measured in **(E)**. **(E)** Strychnine concentration at the soma (blue) and at the OFF-layer dendrites (gray), estimated by the Alexa 594 signal relative to that in the puffer pipette. Timing of light flashes indicated in gold. Signals measured within the ROIs indicated in **(D)**. **(F)** OFF_s_α spike responses to light flashes of varying (0.071, 0.19, 0.52, 1.4 R*/rod) under control conditions (black), immediately following a strychnine puff (red), and following strychnine wash. **(G)** Summary plot (n = 7) showing minimum firing rate, as a percentage of baseline, versus flash intensity for the different conditions described in **(F)**. **(H)** Flash evoked IPSCs, recorded under the same conditions described in **(F)**. **(I)** Summary plot (n = 8) showing IPSC amplitude versus flash intensity for the conditions described in **(F)**.

We sought to block ON-layer inhibitory inputs specifically with 300-ms puffs of the GlyR antagonist strychnine (2-5 μM) locally in the GCL just prior to the light stimulus (Figure 6C-I). Alexa 594 was included in the puffer pipette, and we imaged its diffusion near the recorded OFF_s_α soma and in the OFF layer of the IPL (Figure 6D). Strychnine concentration was estimated by comparing the Alexa 594 fluorescence signal with that generated by the known concentration within the pipette. During the 100-ms period following the light stimulus, estimated [strychnine] exceeded 1 μM at the GC soma, but in the OFF layer it remained below its IC_50_ for GlyRα_1_ (36 nM; Grudzinska et al., 2005; Figure 6E). Strychnine application elicited a small (though statistically insignificant) increase in baseline firing rate (from 42.9 ± 15.0 Hz to 53.4 ± 20.3 Hz, n = 7 cell-attached recordings, *p* = .067, paired *t*-test), so we calculated the light-evoked reduction in firing rate as a percentage of the pre-stimulus baseline rate. Strychnine reduced this OFF response over a range of flash intensities (Figure 6F,G). In whole-cell recordings, strychnine applied in the GCL reversibly reduced flash-evoked IPSCs by an average of 47% across all stimulus intensities (n = 8; Figure 6H,I), suggesting that ON/GCL inhibitory inputs influence scotopic visual responses in OFF_s_α GCs.

### OFFα GC inhibitory RFs are smaller than their dendritic fields

Excitatory synaptic inputs to GCs typically are distributed evenly across the dendritic arbor (Figure S1; Bleckert et al., 2013), so that GC RFs reflect dendritic field (DF) dimensions (Brown and Major, 1966; Boycott and Wassle, 1974; Cleland et al., 1975). RF size typically is measured by delivering light stimuli to different regions of the RF and fitting the spatial distribution of responses with a Gassian function (Enroth-Cugell et al., 1983, but see Rhoades et al., 2019). Strong RF-DF correlations require that the strength of synaptic inputs is relatively even across the dendritic arbor. Inhibitory RFs are usually broader than excitatory RFs (Kuffler, 1953), because the lateral extent of inhibitory connectivity is typically greater than that of excitatory bipolar cells. Under scotopic conditions, particularly when visual stimuli are delivered from darkness, excitatory inputs to OFFα GCs are weak and evoked input is predominantly inhibitory (Murphy and Rieke, 2006; Margolis and Detwiler, 2007; Murphy and Rieke, 2008; van Wyk et al., 2009). We postulated that OFFα GC inhibitory RFs would be smaller than their DF if they were dominated by input from central A2s.

To test this, we recorded light-evoked postsynaptic currents under scotopic conditions from ONα and OFFα GCs and measured RF dimensions (Figure 7). Vertical and horizontal bars of light (0.5 R*/rod/s) were presented at different *x* and *y* displacements, respectively and responses in each spatial dimension were fit with a Gaussian function (Figure 7A; see STAR Methods); the resulting space constants (λ_x_ and λ_y_) were averaged together (λ_RF_). For comparison with anatomical dimensions, RF diameter was calculated from the region over which the function remained >5% of peak (2√3* λ_RF_; Figure 7D).

**Figure 7.**
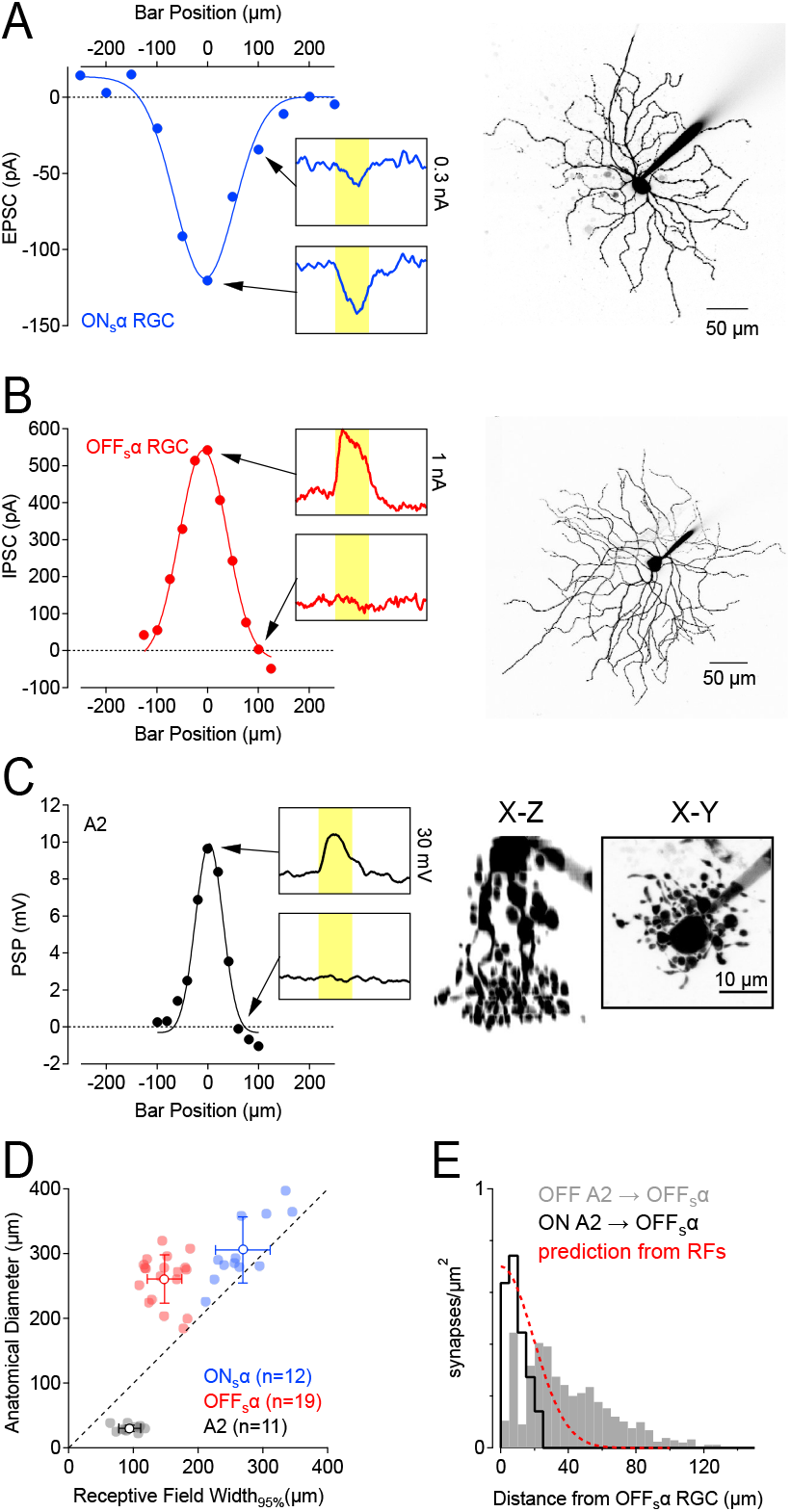
A2 input condenses OFFα GC RFs. **(A)** EPSCs recorded from an ON_s_α GC in response to bars of light presented to different regions of its RF. *Right*, fluorescence micrograph of recorded cell. **(B)** IPSCs recorded from an OFF_s_α GC in response to bars of light presented to different regions of its RF. *Right*, fluorescence micrograph of recorded cell. Also see Figure S4, which shows that blocking surround inhibition with TTX does not influence these RF measurements. **(C)** EPSCs recorded from an A2 AC in response to bars of light presented to different regions of its RF. *Right*, fluorescence micrograph (side and top views) of recorded cell. **(D)** Summary plot comparing DF diameter and RF width in ON_s_α GCs, OFF_s_α GCs and A2 ACs. **(E)** Histogram showing the spatial density distribution of A2→ OFF_s_α inputs in the OFF later (gray) and ON layer (black). Red dashed line indicates predicted spatial distribution of A2 inputs to OFF_s_α GCs based on the relative size of their RFs.

To measure the DF diameter in the same experiments, cells were filled with Alexa 488 during whole-cell recording and imaged after the experiment (Figure 7A, *right*); a convex polygon drawn around the dendritic arbor was used to calculate the average diameter of the DF. In ON_s_α GCs, the excitatory RF diameter was highly correlated with DF diameter (*r* = .84, n = 12, *p* = .00059; Figure 7A,D). OFF_s_α GCs, by contrast, exhibited inhibitory RFs that were narrower than (n = 19, *p* = 6×10^−9^, paired *t* test) and poorly correlated with their DFs (*r* = −.11, *p* = .65; Figure 7B,D). As a result, OFF_s_α RF area constituted a smaller fraction of its DF area when compared to ON_s_α GCs (p = 1.6×10^−5^, Mann-Whitney test). RF dimensions were not altered significantly by TTX (RF diameter in TTX: 95 ± 11% of control, n = 8 OFFα GCs, *p* = 0.19, paired *t* test; Figure S3), indicating that RFs measured this way were not influenced by surround inhibition from wide-field, spiking ACs (Park et al., 2020). When stronger stimuli were used to evoke reliably detected EPSCs, the inhibitory RF diameter was 27 ± 15% smaller than the excitatory RF in OFF_s_α GCs (n = 7, *p* = .015, paired *t* test).

A2 RFs, measured by evoking PSPs with 0.5 R*/rod/s stimuli, were wider than (p = 4.3×10^−7^, paired *t* test) and poorly correlated with their DFs (*r* = −0.15, p = .65; Figure 7C,D). A2 RFs were narrower than OFF_s_α RFs (p = 2.3×10^−7^, *t* test; Figure 7D), suggesting that OFF_s_α GCs are inherited from multiple A2s. The distribution of A2 inputs required to generate the OFF_s_α inhibitory RF was estimated by deconvolving the average A2 RF Gaussian from that of the OFF_s_α inhibitory RF (red line, Figure 7E). This prediction most closely matched the spatial distribution of ON layer A2→OFF_s_α synapses (black line, Figure 7E), suggesting that perisomatic inputs from A2 heavily influence OFFα GC inhibitory RFs.

## Discussion

The results presented here reveal an unexpectedly direct route for visual signals through the rod pathway of the mammalian retina. Anatomical and physiological data show that a small minority of A2s inhibit the somatic region of OFFα GCs, conferring high visual sensitivity and small inhibitory RFs. The close proximity of RB inputs to these outputs within A2 arboreal dendrites ensures efficient signal transfer from single RBs to OFFα GCs. This previously unrecognized motif breaks three established “rules” of retinal circuitry: 1) it violates the laminar segregation of inputs and outputs within narrow-field (NF) ACs mediating “crossover inhibition” in the IPL; 2) it generates RFs that are poorly predicted by dendritic dimensions; 3) it reveals heterogeneous synaptic organization within a single cell type.

### A2s “cross over” within the ON layer

Segregated ON and OFF signaling in the IPL is a consistent hallmark of vertebrate retinas (Famiglietti and Kolb, 1976). ON and OFF BPs provide excitatory input to ON and OFF GCs in the inner and outer IPL, respectively, creating a strong correlation between morphology and physiology (e.g., Wu et al., 2000; Wässle et al., 2009; Baden et al., 2016; Franke et al., 2017, but see Zhang et al., 2007; Hoshi et al., 2009). Wide-field (WF, typically GABAergic) and NF (glycinergic) ACs influence this circuitry in different ways: WF ACs typically mediate lateral interactions within IPL strata, conferring surround feedback or feedforward inhibition to BP terminals or GC dendrites, respectively (Vaney, 1990; Franke et al., 2017), and also inhibiting other ACs (Eggers and Lukasiewicz, 2010). NF ACs operate more vertically, relaying signals between IPL layers – “crossover” inhibition (Roska and Werblin, 2001) that typically influences feedforward signals locally via “push-pull” interactions with excitatory input (McGuire et al., 1984; van Wyk et al., 2009; Franke et al., 2017). NF ACs, including A2s, are thought to reinforce the ON-OFF segregation by providing ON-driven inhibition to the OFF layer, or vice versa (Famiglietti and Kolb, 1975). We show here that A2s contradict this convention, providing powerful feedforward inhibition directly onto OFFα GC somas and proximal dendrites in the ON layer. Future work may reveal whether A2s represent an exception among NF ACs or instead exemplify a previously unrecognized motif of retinal circuitry.

### A2 ON/GCL inputs exert a disproportionate influence on OFFα RFs

Consistent with previous reports (Bleckert et al., 2013), we found that synaptic inputs to *OFFα* GCs are distributed evenly across the dendritic arbor (Figure 1, S1), an arrangement that typically ensures a close agreement between DF and RF dimensions (e.g., Yang and Masland, 1994). Our physiological experiments revealed, however, that A2 input to the proximal dendrites, soma and axon dominates OFF_s_α inhibitory RFs, causing them to be smaller than expected from the breadth of inhibitory inputs across the dendritic arbor (Figure 7D). Each A2 ON/GCL input could provide a larger inhibitory conductance than A2 synapses in the OFF layer, and/or ON/GCL inputs could exert more influence by virtue of their electrotonic proximity to the soma. We favor the latter mechanism, which could accentuate the experimentally assessed contribution of ON/GCL inputs owing to imperfect space clamp and, more physiologically, to light-evoked changes in spiking initiated at the axon initial segment.

### Does the mouse retina contain multiple A2 subtypes?

Neuronal cell types typically are identified based on physiological characteristics, morphological features, synaptic connectivity and gene expression. In the retina, physiological and molecular classifications of bipolar and ganglion cell types agree remarkably well (Baden et al., 2016; Shekhar et al., 2016; Franke et al., 2017; Laboissonniere et al., 2019; Tran et al., 2019), and cells of one type exhibit consistent connectivity patterns with other cell types (Helmstaedter et al., 2013). One previous study divided A2s into two subtypes based on visual sensitivity (Pang et al., 2012), suggesting that A2s making ON/GCL inputs to OFFα GCs may represent a distinct cell type. Two observations suggest that the different A2s described here do not correspond to those proposed subtypes. First, our EM observations provided no evidence that ON/GCL-projecting A2s would exhibit distinct light sensitivity: A2 making ON/GCL synapses received similar RB inputs compared to those that did not. Second, whereas the previously proposed subtypes constituted roughly equal proportions (55%/45%) of the A2 population (Pang et al., 2012), only 14 of 127 (11%) of the A2s studied in the SBFSEM data set made ON/GCL chemical synapses.

Reliable connectivity between cell types requires that their processes overlap in space and, in many cases, that cells express specific cell-adhesion molecules (Hynes and Lander, 1992; Yamagata et al., 2002). A2 arborial dendrites appear to target OFFα GC somas selectively amidst many other options, suggesting that molecular cell-cell interactions may dictate specific connectivity. A distinct subtype of A2s may express a particular adhesion molecule recognized by OFFα GCs; alternatively, all A2s could be molecularly similar but only those located above OFFα GC somas are afforded the geometric opportunity to make ON/GCL contacts. A2s appear to constitute a single cell type with respect to gene transcription (Yan et al., 2020), favoring the second possibility, but this remains an interesting question for future research.

### Relevance to visual signaling in OFFα GCs

Our results highlight a more direct pathway for scotopic signals to reach OFF GCs. A2 ON/GCL outputs are located near RB inputs, ensuring efficient transfer of synaptic signals (Figure 3), and the inhibitory inputs to OFFα GC somas strongly influence light-evoked spike responses (Figure 6). These circuit features may underlie the high sensitivity of OFF GCs (Ala-Laurila and Rieke, 2014), but specific roles for this pathway in night vision remain to be determined. A recent report suggests that some visually guided behaviors in scotopic conditions rely primarily on ON pathway signals that sacrifice absolute sensitivity to reduce noise (Smeds et al., 2019). The OFF pathway that we describe may confer greater sensitivity to high-frequency scotopic signals, reflecting fundamentally asymmetric signaling tasks in the ON and OFF pathway when photons are scarce (Pandarinath et al., 2010).

At higher light levels, A2s relay electrical synaptic input from ON CBs to OFF GCs, contributing to “push-pull” RF subunits (McGuire et al., 1984; Roska and Werblin, 2001) that render OFF_t_α GCs sensitive to looming, dark stimuli (Münch, et al., 2009). This function may require dendritic A2 inputs spread across retinotopic space, rather than the strong somatic input described here, and/or other circuit features (Murphy and Rieke, 2011). ON CBs make gap junction with A2 arboreal dendrites (Strettoi et al., 1992), but we were unable to identify them in this SBFSEM data set to measure their proximity to synaptic outputs to OFFα GCs. Further study is required to identify roles for A2 ON/GCL inputs in OFFα GC signaling under different visual conditions.

## Acknowldgements

This work was supported by the NINDS Intramural Research Program (NS003039 to JSD), the National Eye Institute (EY017836 to JHS, EY028111 to FR) and an unrestricted grant from Research to Prevent Blindness, Inc. to UW Madison Department of Ophthalmology and Visual Sciences (MH).

## Author Contributions

Conceptualization, WNG, MH, FR, JHS and JSD; Methodology, WNG, JHS, JSD; Investigation, MS, WNG, MH, AN, HT, MH, JHS; Writing – Original Draft, JSD; Writing – Review & Editing, MS, WNG, MH, JHS, JSD; Visualization – WNG, MH, MM, JHS, JSD; Supervision – WNG, JHS, JSD; Funding acquisition, MH, FR, JHS, JSD.

## STAR Methods

### Animal Care

All experiments were performed in accordance with protocols approved by the NINDS Animal Care and Use Committee (ASP-1344). Animals were maintained on a 12:12 light-dark cycle and provided free access to food and water. Prior to electrophysiological experiments or immunohistochemistry, mice were deeply anaesthetized with isoflurane (Baxter) and euthanized via decapitation.

### SBFSEM Analysis

A 3D SBFSEM data set (k0725) of a 50×210×260 μm^3^ block of P30 C57BL/6 mouse retina (13.2×13.2×26 nm^3^ voxels; Ding et al., 2016) was analyzed as described previously (Graydon et al., 2018; Park et al., 2020). A2s were identified based on morphological characteristics and synaptic input from RBs. RB inputs were distinguished by their location deep in the IPL and presynaptic ribbons that are larger than those in CB terminals (Strettoi et al., 1992; Graydon et al., 2018). OFF_s_α and OFF_t_α GCs were identified by soma diameter and dendritic ramification in the distal IPL. Skeletonization and annotation were performed manually using Knossos (Helmstaedter et al., 2011). Voxel coordinates were tilt-corrected and normalized either to to the positions of the ON and OFF SACs (Ding et al., 2016).

### Immunohistochemistry (Figure 2A-H)

Whole retinas were dissected from C57BL6-WT mice (^~^P60) and fixed in 4% paraformaldehyde (PFA) in 0.1M phosphate buffer solution (PBS) for 20 min at room temperature (RT), then rinsed with PBS. Retinas were pre-incubated for 5 h at RT or overnight at 4°C in blocking solution containing 10% donkey serum (Sigma, MO), 2% BSA, 0.5% Triton X-100 and 0.1% sodium azide in PBS. Retinas were then incubated for three nights at 4°C with primary antibodies against RBPMS (guinea pig polyclonal, 1:500; Phosphosolution), GlyRα_1_ (rabbit monoclonal, 1:500; Synaptic Systems) and SMI32 (mouse monoclonal, 1:500; Millipore-Sigma) in blocking solution. Retinas were then rinsed 3 times in PBS, incubated overnight at 4°C with secondary antibodies against guinea pig (donkey, Alexa Fluor 488, 1:500; Invitrogen), rabbit (donkey, Cy3, 1:500; Jackson ImmunoResearch) and mouse (donkey, Cy5, 1:500; Jackson ImmunoResearch) in blocking buffer, rinsed in PBS and thereafter mounted with DAPI mounting medium (Vector Labs). Immunoreactivity was imaged using a confocal microscope (Zeiss LSM 580, 20×/1.0NA objective).

### Single-cell immunolabeling (Figure 2I-L)

GCs in whole-mount retinas were filled with 4% Neurobiotin through a whole-cell patch electrode, then fixed for 15 mins at RT in artificial cerebrospinal fluid containing 4% paraformaldehyde. Post-fixation, retinas were washed in 0.1M PBS, pre-incubated overnight at 4°C in blocking solution containing 5% donkey serum and 0.5% Triton in PBS and then incubated for 3 nights at 4°C with primary antibody against GlyRα_1_ (mouse monoclonal mAb2b, 1:500; Synaptic Systems) in blocking solution. Retinas were subsequently rinsed in PBS, incubated with a secondary antibody against mouse (goat anti-mouse 488, 1:000; Invitrogen) and streptavidin conjugated to Alexa Fluor 568 (1:1000; Invitrogen), rinsed in PBS and mounted using Vectashield anti-fade mounting medium (Vector labs).

Images were acquired using a confocal microscope (Leica SP8, 63×/1.4NA objective). Image stacks encompassing the entire GC (soma and dendritic arbor) were acquired, median-filtered in FIJI (NIH) and visualized with Amira software (Thermo Fisher Scientific). Individual GCs were masked in 3D via the *LabelField* function in Amira. Once a GC was isolated in a 3D mask, the GlyRα_1_ receptor channel was multiplied with the GC mask to isolate the GlyRα_1_ signal within the GC. For quantification of the GlyRα_1_ signal within the ON (soma and dendritic arbor within the ON lamina) and OFF (dendritic arbor within the OFF lamina) arbors of the GC, the volume of GlyRα_1_ signal within the RGC compartment and above background (threshold of 3 standard deviations above the noise peak applied to eliminate background pixels) was estimated and normalized to the GC compartment volume (see also Hoon et al., 2017).

### Live tissue preparation and electrophysiology

Eyes were removed from adult mice (P30-80) of either sex and retinas were isolated at room temperature (ChR2 and Ca^2+^ imaging experiments) or 32-34°C (scotopic RF measurements and somatic puff experiments) in bicarbonate-buffered Ames media (Sigma) equilibrated with carbogen (95% O_2_/5% CO_2_). Tissue was trimmed and mounted GCL up (i.e. whole-mount), placed under a two-photon laser-scanning microscope (Zeiss 510 or Scientifica Hyperscope) and superfused during experiments with Ames (which contains 1.1 mM Ca and 1.2 mM Mg) at 30-34°C (4-8 mL/min). For experiments examining retinal signaling in response to visual stimuli, mice were dark adapted for >2 hrs prior to dissection and retinas were isolated under infrared illumination (940 nm LED light source, Thorlabs). Retinas used for for Ca^2+^ imaging experiments were isolated under dim red light. Cells within the whole-mount retina were selected for recording under differential interference contrast (DIC) optics (940 nm): α GCs were identified by their large soma size, and A2s by their pear-shaped somas that jut into the IPL from the INL.

Cell-attached and whole-cell recordings were made with an electronic amplifier (MultiClamp 700B, Molecular Devices) controlled with custom software written in IgorPro or Matlab (Symphony: https://symphony-das.github.io). Patch electrodes (1.5mm OD borosilicate glass, 4-5 MΩ for GCs, 5.5-6.5 MΩ for A2s) were filled with internal solution (280-285 mOsm, pH 7.4) containing (mM): 90 CsCH_3_SO_3_ or KCH_3_SO_3_, 20 TEA-Cl, 4 MgATP, 0.4 NaGTP, 10 EGTA, 10 Na_2_ Phosphocreatine, 10 HEPES.

### Ca^2+^ indicator imaging

Ca^2+^ indicator imaging experiments were performed on a Zeiss (Thornwood, NY) LSM-510 confocal microscope (40x/1.0 DIC W Plan-Apochromat objective) controlled by ZEN 2009 software and custom macros (Igor Pro, WaveMetrics, Lake Oswego, OR). A2s were patched in whole mounts and filled with a Cs^+^-based intracellular solution containing Alexa 594 hydrazide (50 μM; Thermo Fisher Scientific) to visualize cell morphology and the Ca^2+^ indicator Fluo-5F (150 μM; Thermo Fisher Scientific). No additional intracellular Ca^2+^ buffers were present during imaging experiments. Imaging commenced 10-15 minutes after establishing whole-cell recordings to allow dyes to diffuse throughout the cell. Alexa 594 and Fluo-5F were excited by 543-nm He-Ne and 488-nm Ar lasers. Transient indicator signals were evoked by 100 ms voltage steps applied through the recording pipette every 15-20 seconds, measured in (bidirectional) frame scan mode (10.4 Hz) and low-pass filtered (2-5 Hz) offline. Glutamatergic transmission and light responses were routinely blocked pharmacologically by DNQX (20 μM; Abcam), R-CPP (5 μM; Tocris Bioscience) and L-AP4 (20 μM, Abcam) before imaging started. Image analysis was performed using custom macros in IgorPro. Individual lobular and arboreal varicosities were analyzed as single regions of interest (ROIs) and all data are presented as ΔF/F. Z-stacks (1 μm intervals, 4 images averaged at each z level) were acquired at the end of each experiment to reconstruct cell morphology.

### Optogenetic stimulation

Optogenetic stimulation experiments were performed on Zeiss LSM-510 (40×/1.0 NA objective) and a Nikon C1 (40×/0.8 NA objective). *Neurod6^Cre^* and Ai27 (Gt(ROSA)26Sor) lines were crossed to yield mice that expressed ChR2 and tdTomato in A2s and at least one other narrow-field AC type. A2s were distinguished from other labelled amacrines due to their stronger tdTomato expression and their arboreal dendrites that extended through the entirety of the IPL and sometimes into the GCL (the dendrites of other labelled ACs were constrained to IPL sublaminas 1-4). ChR2 was activated with a 500 ms spot scan from a 488 nm Ar laser centered over an AC soma. Stimulation timing was verified by diverting a fraction of the laser light to a photodiode. ChR2-evoked IPSCs were recorded from GCs in whole-cell voltage clamp (V_hold_ = 10 mV) with a Cs-based internal solution supplemented with Alexa488 (50 uM). Photoreceptor-driven light responses to the 488 nm laser were eliminated by blocking all glutamatergic synaptic transmission in the retina (e.g. eliminating bipolar transmission) with L-AP4 (20 μM; Abcam), UBP 310 (25 μM; Abcam), NBQX/DNQX (10 μM/20 μM; Abcam) and D-AP5/R-CPP (25 μM/5 μM; Tocris Bioscience). A2-GC distance was determined by collecting a composite z-stack and measuring the linear distance between the centers of the A2 (red) and GC (yellow) somas in the *x-y* plane.

### Visual stimulation

Visual stimuli were controlled by Symphony, generated with a customized DMD projector (4500 Light Crafter, National Instruments; Franke et al., 2019) and a 405 nm LED (Thorlabs), attenuated by neutral density filters (^~^6 log units) and routed through the microscope condenser, which was was adjusted so that images were in focus at the level of the rod outer segments. Spectral and power densities were measured at the sample plane and converted to R*/rod/s as previously described (Grimes et al., 2014). Scotopic RFs of A2 amacrine cells and GCs were probed with 50 × 500 μm^2^ bars (250 ms) that elicited 0.1-1 R*/rod. Excitatory RFs of ON_s_α GCs and A2s were probed at 50 and 25 μm intervals, respectively; inhibitory RFs in OFF_s_α GCs were probed at 25 μm intervals. Bars were presented 5-10 times at each location and in two orthogonal directions. Responses to bars at each location were averaged and baseline subtracted prior to measuring amplitude by averaging over a window 100-350 ms following stimulus onset. Response amplitudes were plotted versus bar position and fit by a sum of two Gaussians (one for the center and a second, larger one of opposite polarity for the surround). Surround amplitudes typically were small relative to those in the center.

### Puff application of strychnine

Contributions from somatic glycine receptors were examined by locally puffing strychnine onto the GC soma while recording responses to small (50 μm diameter) spots (17 ms flash) positioned in the center of the GC’s RF. Puff electrodes (6-8 MΩ) were filled with 2-10 uM strychnine (to block glycine receptors) and 50 uM Alexa594 (to monitor the diffusion of the puff and estimate strychnine concentration over space and time). Small holes were torn into the inner limiting membrane on either side of the GC somas to create room for the 2 electrodes. Puffs were applied (Picospritzer, Parker) for 300 ms prior to the flash; trials were repeated at 5-6 s intervals. Flashes of four different intensities were presented, and average responses (5-7 trials) were obtained for each flash strength in each condition (control, puff, wash). Data from the first 8 flashes in each condition were excluded from further analysis to allow for adaptation to the mechanical puff. Average responses were smoothed (10 ms box filter) prior to measuring minimum spike rate or maximum INH current. Spike suppression was calculated by dividing response minimums by the baseline firing rate. Strychnine concentrations were estimated by measuring average fluorescence within a ^~^15 um^2^ ROI and normalizing this to the maximum fluorescence signal (near the pipette tip or within the pipette).

### Statistical Analysis

Except where noted, normally distributed (as determined by the Jarque-Bera test) group data values are reported as mean ± SD and compared with a *t*-test. Otherwise, data are compared with a Wilcoxon rank test. Pearson’s correlation coefficient (*r*) was calculated with a linear correlation test (Igor Pro). Significant differences were concluded if *p* < .05.

## Supplemental Information

**Figure S1 |** Excitatory (ribbon synapse) inputs to OFF_s_α and OFF_t_α GCs in SBFSEM data set

**(A-B)** Skeletonized OFF_s_α **(A)** and OFF_t_α **(B)** GCs with excitatory inputs annotated (blue spheres). Scale bar in **(A)** applies to both panels.

**Figure S2 |** Other amacrine cell inputs to the OFF_s_α GC

Skeletonization of inputs to ON dendrites reveals several cell types other than A2s that provide synaptic inhibition. **(A)** *i*, a transverse view showing bistratified cells with dendrites ramifying in sublaminae 1 and 5 (S1 and S5) that provide both ON and OFF input to the GC. 8 presynaptic cells in orange make synapses (magenta) onto the GC (blue). *ii*, individual cells are rendered in different colors. *iii*, an en face view (GCL at surface) illustrates heterogeneity in the 8 presynaptic cells; e.g., cell *a* has a larger dendritic field than cell *b*. **(B)** As in **(A)**, illustrating 3 cells with assymetric dendritic fields. Interestingly, all 3 are oriented in the same direction (the retinal location and orientation of the sample is unknown). **(C)** 2 displaced (*i*), 1 S3, S5 bistratified (*ii*), and 3 wide-field (*iii*; not traced back to a soma) cells that make ON synapses with the GC. **(D)** The GC receives input from wide-field cells in multiple sublaminae (*i*); these are shown en face (*ii*), as in **(A***iii*).

**Figure S3** | Surround inhibition from spiking amacrine cells does not affect OFFα GC RF size

**(A)** Amplitudes of IPSCs (V_hold_ = + 10 mV) evoked in an OFF_s_α GC with light bars located at various distances from the center of the GC’s RF (see STAR Methods) in control solutions (black) and in the presence of 0.5 μM TTX (blue). Solid lines indicate Gaussian fits. **(B)** IPSCs recorded in the same cell shown in **(A)**, evoked by bars at the positions indicated. **(C)** Summary graph comparing RF width in control and TTX for 4 OFF_s_α GCs and 4 OFF_t_α GCs.

