## Supplementary figures and images for "Dendro-somatic synaptic inputs to ganglion cells violate receptive field and connectivity rules in the mammalian retina"

### Supplemental Figure

#### Figure S1

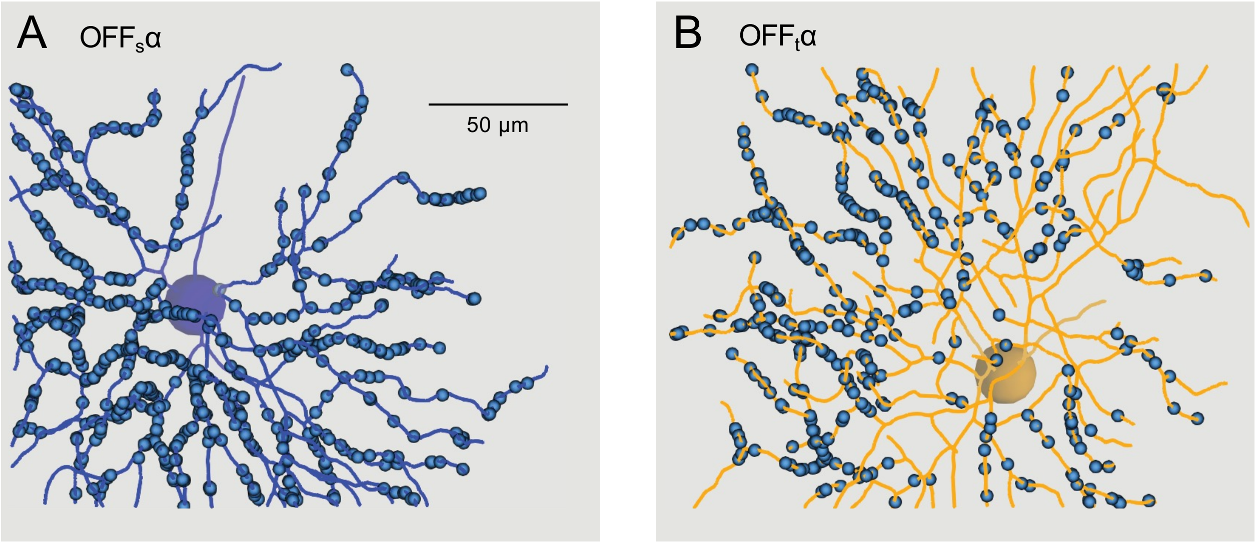

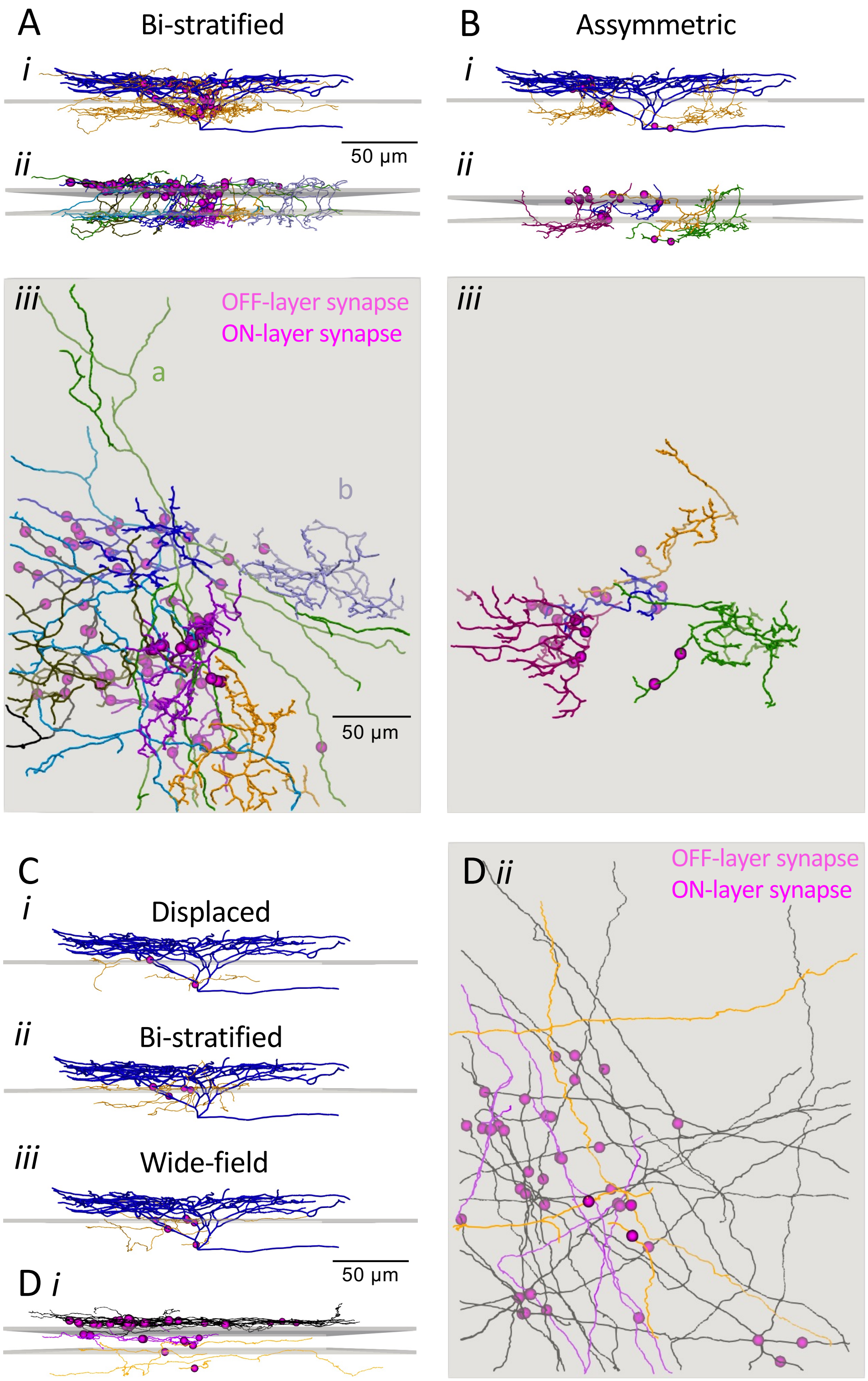

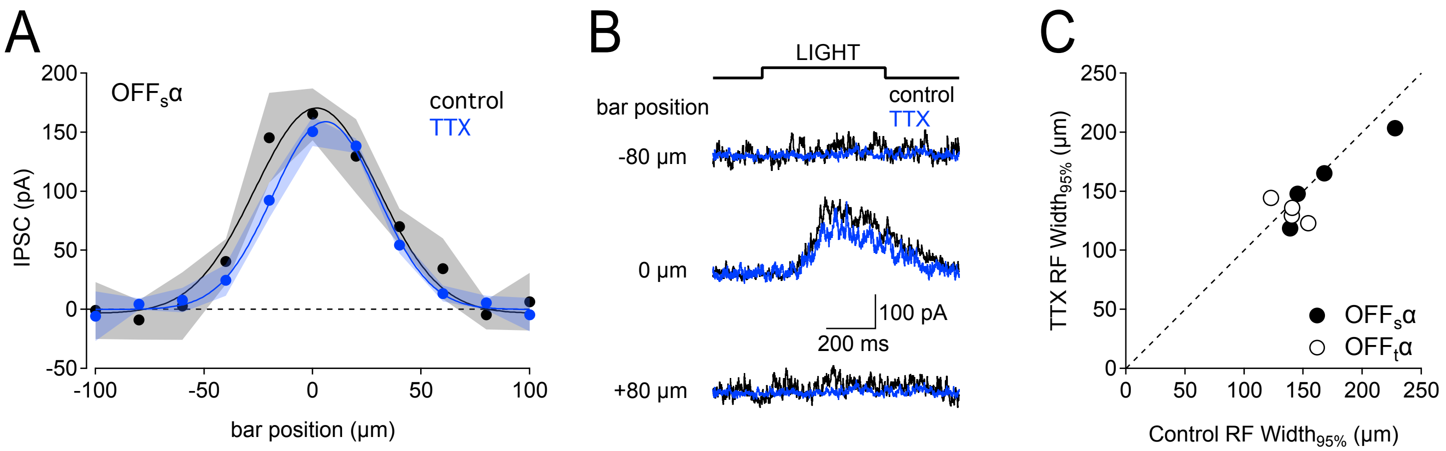
